# The signalling axis CDK12-BRCA1 mediates dinaciclib associated radiosensitivity through p53-mediated cellular senescence

**DOI:** 10.1101/2024.04.25.590582

**Authors:** Natalia García Flores, Diego M. Fernández-Aroca, Cristina Garnés-García, Andrés Domínguez-Calvo, Sebastia Sabater, Ignacio Andrés, Jaime Jiménez-Suárez, Pablo Fernández-Aroca, Francisco Cimas, Guillermo de Cárcer, Borja Belandia, Ignacio Palmero, Pablo Huertas, María José Ruiz-Hidalgo, Ricardo Sánchez Prieto

**Author notes:** Correspondence should be addressed to: Ricardo Sánchez Prieto, Ph.D.

## Abstract

Pan-CDK inhibitors are a new class of targeted therapies that can act on multiple CDKs, with dinaciclib being one of the most promising compounds. Although used as monotherapy, an interesting approach could be to combine it with radiotherapy. Here we show that dinaciclib increases radiosensitivity in some experimental models of lung and colon cancer (A549 or HCT 116) but not in others (H1299 or HT-29). Dinaciclib did not alter responses such as ATM signalling, apoptosis or cell cycle profiling after ionising radiation exposure that have been described for other CDK inhibitors. Mechanistically, inhibition of CDK12 by dinaciclib abolishes BRCA1 expression, which blocks homologous recombination (HR), leaving only the non-homologous end joining repair process (NHEJ), which ultimately promotes the induction of ionising radiation-associated cellular senescence in a p53-dependent manner, explaining the lack of effect observed in some experimental models. In conclusion, our report explains the molecular mechanism involved in this novel therapeutic effect of dinaciclib and provides a rationale for more selective and personalised chemo/radiotherapy treatment according to the genetic background of the tumour.

**Highlights:**

➢ Dinaciclib is a radiosensitising agent in lung and colon cancer cell lines.
➢ Dinaciclib promotes radiosensitivity by inhibiting CDK12 leading to a decrease in BRCA1 expression.
➢ Lack of BRCA1 blocks homologous recombination response to DNA damage, enhancing ionising radiation-associated senescence in a p53-dependent fashion.
➢ The combination of dinaciclib plus radiotherapy could be a novel treatment for tumours holding functional BRCA1 and p53 signalling.

## 1.1. Introduction

The cell cycle has become one of the major targets in cancer therapy. Indeed, in recent years there have been significant advances in targeting the cellular machinery of the cell cycle, with a particular focus on cyclin-dependent kinases (CDKs) (for a review see [1]). In fact, CDK4/6 inhibitors are currently used in clinical practice to treat metastatic ER+ breast cancer [1] and are also being considered for use in other cancers [2]. In addition to CDK4/6 inhibitors, a new class of CDK inhibitors has been proposed with a broader spectrum of activity, acting on multiple CDKs simultaneously, of which dinaciclib is a paradigmatic example. This novel pan-CDK inhibitor has a preferential effect on CDK1, 2, 5 and 9 with a lower affinity for CDK4 and CDK6 [3]. In addition, recent evidence has shown that CDK12 is a novel target for dinaciclib [4], increasing its potential in cancer therapy [5]. In fact, clinical trials are underway to evaluate the use of dinaciclib in various tumour types [6,7]. In addition, to a clear anti-tumour effect in various experimental models, from osteosarcoma [8] to cholangiocarcinoma [9], dinaciclib has shown an exceptional ability to enhance the efficacy of conventional chemotherapy [10–12] or novel therapeutic approaches [13]. However, no reports have demonstrated the molecular basis for the potential use of dinaciclib in combination with radiotherapy. Interestingly, there is increasing evidence linking dinaciclib to the DNA repair machinery through effects on molecules such as BRCA1 [4,10,14].

With this in mind, we decided to evaluate the potential of dinaciclib as a radiosensitising agent. Our data show that dinaciclib exerts a potent radiosensitising effect in various experimental models of colon and lung cancer by promoting ionising radiation (IR)-dependent senescence. This effect is the consequence of CDK12 inhibition leading to downregulation of BRCA1 block HR, which triggers p53-dependent IR-associated senescence. In conclusion, our data demonstrate the potential of dinaciclib in combination with radiotherapy and describe the signalling axis that mediates this effect, allowing the correct selection of patients who could benefit from this therapeutic combination.

## 2. Material and methods

### 2.1. Cell lines, plasmids, chemicals and antibodies

A549 and H1299 (lung cancer), HCT 116 and HT-29 (colon cancer), HEK293T and U2OS cells have been cultured as previously described [15]. Cells were maintained in 5% CO_2_ and 37°C; and grown in Dulbecco’s modified Eagle’s medium supplemented with 10% fetal bovine serum, 1% glutamine and 1% Penicillin/Streptomycin. All cell culture reagents were provided by Lonza.

Plasmids used for short hairpin RNA (shRNA) interference were as follows: Human PLKO.1-puro-shRNAp53 and Empty vector has been previously described [15]. pLKO.1-puro-shRNACDK12 (TRCN0000196423 and TRCN0000368335) and pLKO.1-puro-shRNABRCA1 (TRCN0000244984 and TRCN0000244987) were purchased from Merck.

Dinaciclib (MedChemExpress), navitoclax (MedChemExpress) and ATM inhibitor Ku-55933 (Calbiochem) were dissolved in DMSO, aliquoted and stored at −80 °C. Puromycin (Sigma-Aldrich) were dissolved in double-distilled water, aliquoted and stored at −20 °C.

Antibodies used for western blot or immunofluorescence are summarized in supplementary table 1.

### 2.2. Transfections and infections

Lentiviral production and cell infection were performed as previously described [15]. In all knockdown experiments, infected cells were used one week after selection with puromycin. Each experiment was performed with 3 different pools of infection. Infected cells were maintained for a maximum of 2 weeks after selection and then discarded and replaced with a new fresh pool of infected cells.

### 2.3. Western Blotting

Cells were collected in RIPA lysis buffer (50 mM Tris, pH 8; 1.5 mM MgCl2; 1 mM EDTA; 1 mM EGTA; 1% Triton X-100; 150 mM NaCl; 20 mM β-Glycerophosphate; 0.1% SDS; 0.5% desoxycholic acid). Protease and phosphatase inhibitors (Merck) were added prior to lysis. Protein quantification and western blotting was performed as previously described [16]. 100 μg, otherwise is indicated, of protein were loaded onto appropriate percentage SDS-PAGE, transferred to PVDF membranes using semi-dry Pierce Power Blot (ThermoFisher) and blotted against different proteins via specific antibodies. Antibodies were detected by enhanced chemiluminescence (Amersham) in a LAS-3000 system (FujiFilm).

### 2.4. Immunocytochemistry

Cells were grown onto SPL cell culture slides (Labclinic) prior to IR. After treatment cells were fixed, permeabilized and incubated with the indicated antibodies as previously described [17]. Positive immuno-fluorescence was detected using a Zeiss Apotome fluorescence microscope and processed using Zen 2009 Light Edition program (Zeiss). Foci quantification was performed with CellProfiler (Broad) [18]. Images show a representative cell from a minimum of 100 quantified (5 fields per sample captured). Data shown are the average of, at least, three independent experiments.

### 2.5. RNA isolation, reverse transcription and Real-time Quantitative PCR

Total RNA obtention, cDNA synthesis and Real time PCR was performed previously described [19]. Primers for all target sequences were designed by using NCBI BLAST software and purchased from Merck as DNA oligos. Primer sequences can be found in Supplementary Table 2. Data shown are the average of, at least, three independent experiments performed in triplicate.

### 2.6. Irradiation and clonogenic assays

Cells were irradiated by the technical staff of Radiotherapy Unit at University General Hospital of Albacete, in a Clinac Low Energy 600C linear electron accelerator from Varian (Palo Alto, California, USA) at a dose rate of 600 cGy/min in a radiation field of 40x40 cm. Cells were plated and the following day, cells were treated with 10 nM dinaciclib for 24 hours prior to IR. Then, fresh medium was added, and cells were incubated at 37°C for 10-14 days to allow colony formation. Clonogenic assays were performed and valuated as previously described [19,20]. Values were referred to unirradiated controls, set at 1. SF2Gy was calculated by applying a linear-quadratic model [21]. Data shown are the average of, at least, three independent experiments performed in triplicated cultures.

### 2.7. Senescence-Associated β-Galactosidase activity (SA-β-Gal)

6 days after IR, cells were washed in PBS, fixed for 5 min (room temperature) in 2% formaldehyde, 0.2% glutaraldehyde, washed twice with PBS for 5 minutes, and incubated for 16 hours at 37°C with freshly prepared SA-β-Gal staining solution as previously described [22]. After this, cells were washed twice with PBS for 5 minutes. Images were acquired at 20X and show a representative field out of 5 acquired per sample (minimum of 100 cells quantified per condition). Data shown are the average of three independent experiments.

### 2.8. Flow cytometry

For cell cycle analysis, 10^5^ cells were seeded in 6 cm plates. 24 hours later, cells were exposed to different treatments. Cell cycle was analysed as previously described at indicated times after IR treatment [15]. Apoptosis, usually 48 hours after IR exposure, was detected with Annexin V-FITC (Immunostep) following manufacturer’s instructions. Samples were processed in a MACSQuant Analyzer 10 (Miltenyi Biotec). Data were analysed by using MACSQuant Analyzer 10 software (Miltenyi Biotec). Data shown are the average of, at least, three independent experiments performed.

### 2.9. Dose-response measurements

For cell proliferation measurements, 10^4^ cells/well were seeded in 24-well plates and proliferation was analysed 16 hours and 3 days after treatment by an MTT-based assay as previously described [16]. Data shown are the average of three independent experiments performed in triplicated cultures.

### 2.10. GFP reporter assay

U2OS cells harbouring a single copy of the reporter constructs DR-GFP (HR; [23]) or EJ5-GFP (NHEJ; [24]) were used as previously described [25]. They were grown in a standard medium supplemented with 1 μg/ml. puromycin (Sigma, P8833). Briefly, 60.000 cells were plated in 6-well plates in duplicate. One day after seeding, cells were treated with dinaciclib (10 nM) or mock treated. The next day, each duplicate culture was infected with lentiviral particles containing I-SceI–BFP expression construct at MOI 10 using 8 µg/ml polybrene in 2 ml of DMEM. Then, cells were left to grow for an additional 24 hours before changing the medium for fresh DMEM. 48 hours after viral transduction, cells were trypsinized after a PBS wash and resuspended in 1 ml of PBS. Samples were analysed with a LSRFortessa™ Cell Analyzer (BD) Flow Cytometer.

### 2.11. Statistical analysis

Data are presented as mean ± standard deviation (S.D). Statistical significance was evaluated by Student’s t test or ANOVA using GraphPad Prism v9.0 software. The statistical significance of differences is indicated in figures by asterisks as follows: *p < 0.05, **p < 0.01 and ***p<0.001.

## 3. Results

### 3.1. Dinaciclib is a radiosensitizing agent that does not affect ATM signaling, apoptosis, or cell cycle in response to IR

First, we analysed the toxicity associated to dinaciclib in different experimental model at different doses. As it showed, (Supplementary information 1A), all cell lines showed a similar toxicity being slightly lower in HT-29, but in all cases with an IC50 between 5 to 10 nM. In addition, the distribution of the cell cycle was analyzed after 24 hours of incubation (10 nM), showing alterations of the cell cycle in all cell lines (Figure 1A). Next, we analyzed the combination of a 24 hours preincubation to 10 nM dinaciclib, with a toxicity lower than 20% in any of the experimental models (Supplementary information 1B), and IR by clonogenic assay. As shown in Figure 1B, 24 hours pretreatment with dinaciclib 10 nM enhanced radiosensitivity in A549 and HCT 116 but failed in H1299 and HT-29, showing a correlation with the presence of a wild type (WT) p53.

**Fig. 1.**
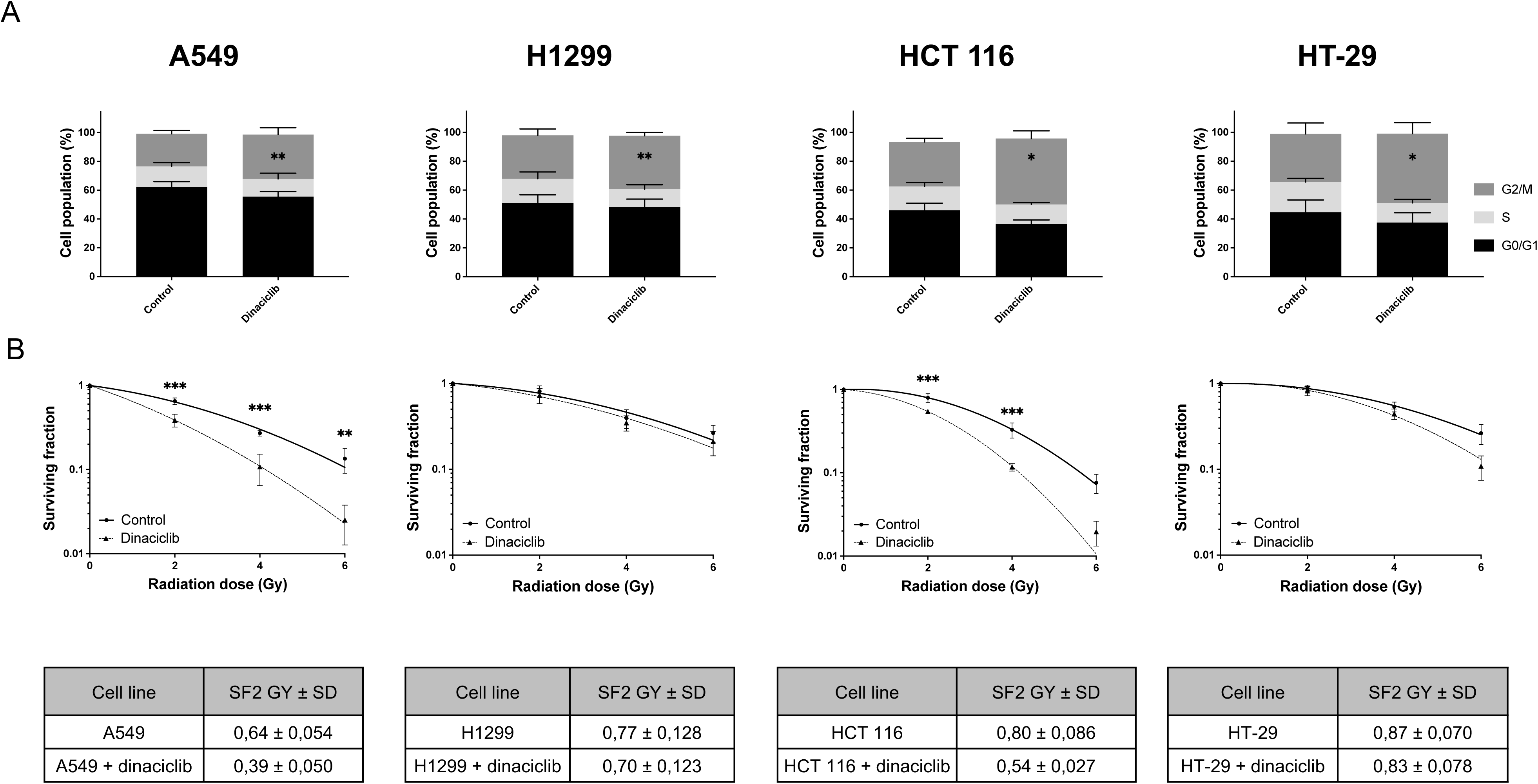
Dinaciclib promotes radiosensitivity in A549 and HCT 116 but not in H1299 or HT-29 cell lines. A) A549, H1299, HCT 116 and HT-29 cell lines were incubated for 24 hours in the presence of vehicle (DMSO) or dinaciclib (10 nM) and the cell cycle was analysed. Histograms represent the average of 3 independent experiment. Bars denote standard deviation (S.D). B) *Upper panels*: Clonogenic assays for A549, H1299, HCT 116 and HT-29. Cells were plated and 24 hours later exposed to vehicle or dinaciclib (10 nM) for additional 24 hours. Then cells were exposed to the indicated doses of X rays and immediately after media was replaced. *Lower panels*: Surviving fraction was normalized to respective unirradiated controls. Curves were fitted using lineal-quadratic model. Bars mean S.D.

In view of our previous findings, we carried out an in-depth analysis of the putative mechanism by which dinaciclib promotes radiosensitivity focusing on two cell lines, A549 and H1299. First, we analysed whether dinaciclib affects ATM activity. Therefore, we irradiated cells in the presence/absence of dinaciclib pretreatment and evaluated ATM signaling. As it shown in Figure 2A, B and C, neither H2AX nor KAP1 phosphorylation after IR exposure were affected by the presence of dinaciclib. Next, we evaluated apoptosis after IR exposure. As shown in Figure 3A and B, 48 hours after IR exposure, dinaciclib did not show a synergistic effect in terms of IR-induced apoptosis, being at best additive, and with identical behavior regardless of the radiosensitising effect. In addition, cell cycle profiles were also evaluated 48 hours after IR. Interestingly, irradiated cells pretreated with dinaciclib, showed a similar pattern to untreated cells (Figure 3C and D), and again this observation was almost identical regardless of the radiosensitizer effect observed.

**Fig. 2.**
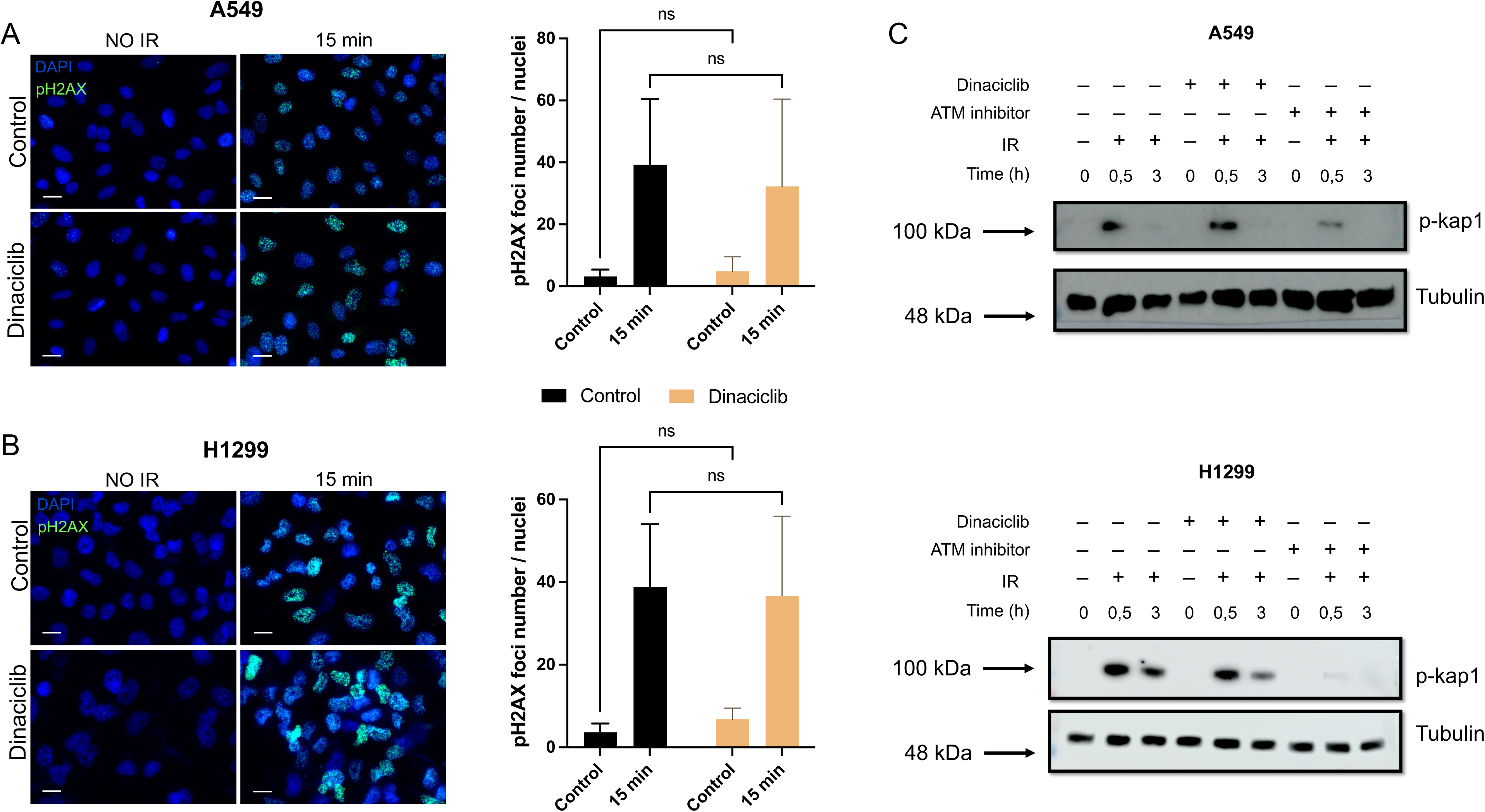
ATM signaling is not targeted by dinaciclib. A) A549 (left panel) cells were plated onto cell culture slides and 24 hours later exposed to vehicle or dinaciclib (10 nM) for additional 24 hours. After irradiation (10 Gy) cells were fixed and processed for immunocytochemistry against phospho-Histone H2AX (Ser139). Images show a representative field out of a minimum of 5 analysed. Scale bars represent 20 μm. Quantification of phospho-Histone H2AX (right panel) foci number per nuclei in three independent experiments. Bars mean S.D. B) Same as in A for H1299 cells. C) KAP1 activation was evaluated in A549 (upper panel) and H1299 (lower panel) cell lines in presence/absence of dinaciclib by western blot at indicated times after exposure to 10 Gy. Cells were treated with vehicle or dinaciclib (10 nM) or ATM inhibitor (10 μM) for 24 hours before IR. Protein extracts were blotted against the indicated antibodies.

**Fig. 3.**
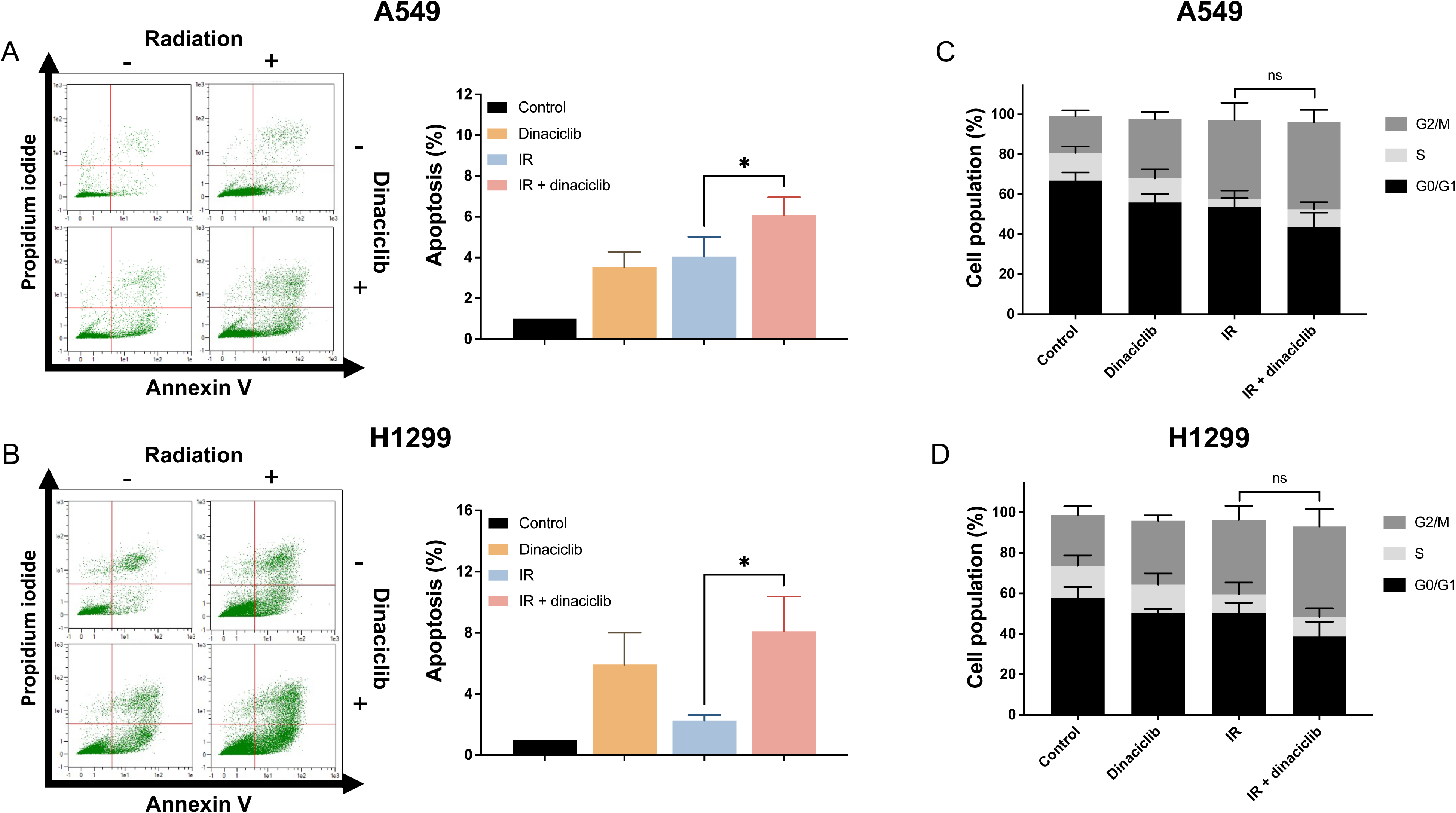
Dinaciclib does not modulates apoptosis or cell cycle in response to ionising radiation. A) *Left panel:* Representative images of apoptosis induction 48 hours after irradiation (10 Gy) in A549 cells pretreated with dinaciclib or vehicle for 24 hours prior to irradiation *Right panel:* Histogram showing the average of three independent experiments. Bars mean S.D. B) Same as in A for H1299 cells. C) Cell cycle was evaluated by flow cytometry 48 hours after IR (10 Gy) in A549 treated as in A. Histogram showing the average of three independent experiments representing the percentage of population in the different phases of the cell cycle. Bars mean S.D. D) Same as in C for H1299 cells.

In conclusion, this series of experiments discards ATM signaling, apoptosis or gross cell cycle alterations as the putative mechanisms of dinaciclib-associated radiosensitivity in our experimental model.

### 3.2. Dinaciclib promotes senescence associated to IR in a p53 dependent fashion

All of the above suggests that other events may be involved in our initial observation. Therefore, we decided to investigate IR-associated senescence. Interestingly, SA-β-Gal staining in the A549 cell line 6 days after IR exposure was significantly increased in the presence of dinaciclib pretreatment compared to separate treatments (Figure 4A and C), showing a synergistic effect (>2.5-fold increase). However, SA-β-Gal staining was barely detected in H1299 cell line (Figure 4B and C), suggesting a possible mechanism of radiosensitivity. Indeed, similar results were obtained in the HCT 116 and HT-29 cell lines (Supplementary information 2). To support this observation, we analysed several genes associated with the Senescence-Associated Secretory Phenotype (SASP) by RT-qPCR, showing an enhanced expression in the A459 cells but not in H1299 cells, a result fully consistent with SA-β-Gal staining (Figure 4D). Next, we decided to use navitoclax, a known senolytic compound [25]. Based on our previous experiments, we incubated cells with navitoclax (1 μM) for 48 hours, 5 days after IR exposure. Interestingly, navitoclax was able to promote a marked increase in radiosensitivity associated with dinaciclib pretreatment in our experimental model of A549 but not in H1299 cells (Figure 4E and F). Indeed, in this experimental setting, SA-β-Gal staining was performed showing a fully concordant result with the senolytic activity of navitoclax (Supplementary information 3).

**Fig. 4.**
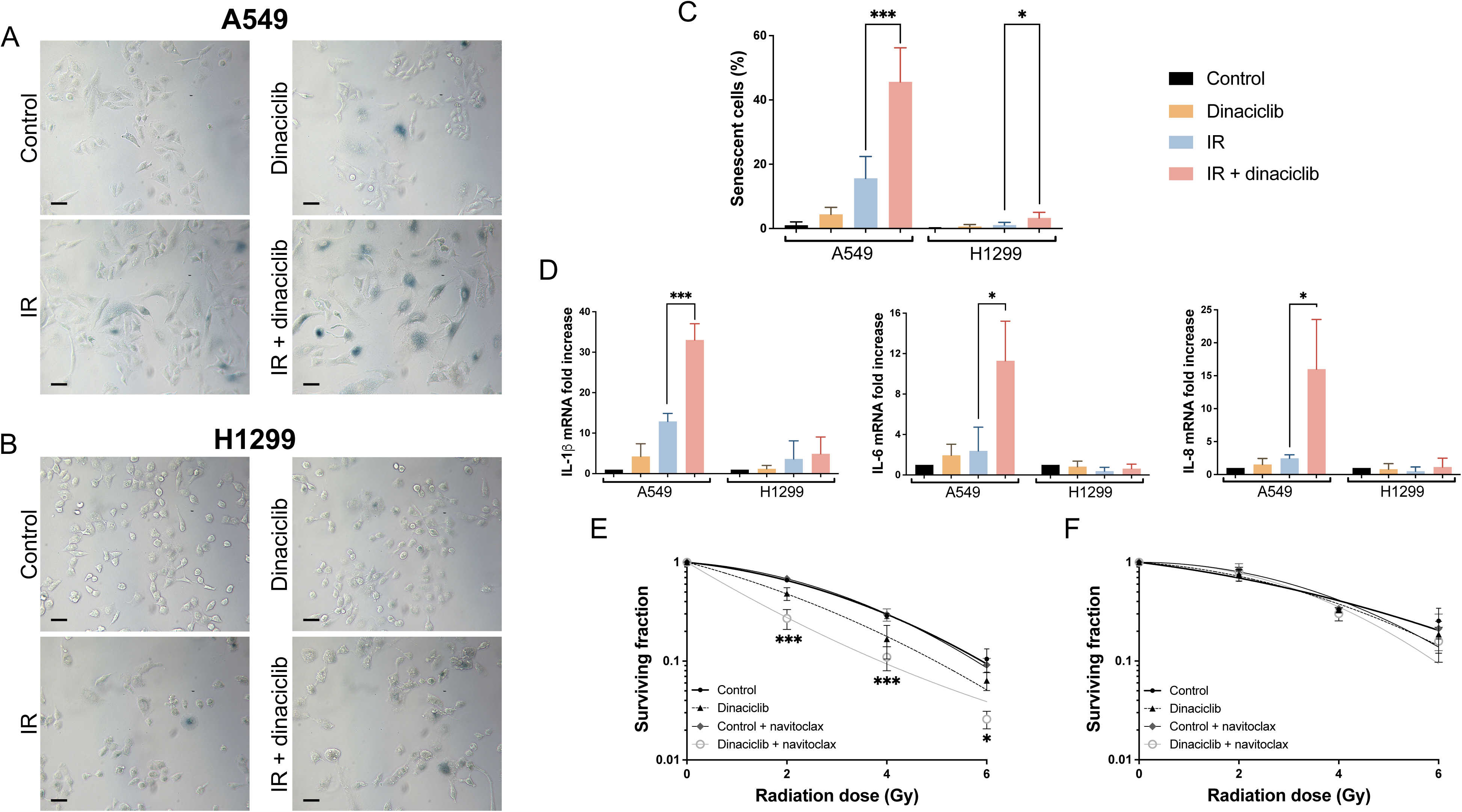
Dinaciclib enhances cell senescence in response to ionising radiation. A) A549 cells pretreated with dinaciclib or vehicle for 24 hours were irradiated (6 Gy) and 6 days later SA-β-Gal activity was detected by X-Gal staining. A representative image is shown. Scale bars represent 100 μm. B) Same as in A) for H1299 cells. C) Histogram showing the average of, at least, three independent experiments representing the percentage of positive senescent cells in A549 and H1299 cell lines. Bars mean S.D. D) Gene expression of indicated SASP genes in different conditions was evaluated 6 days after IR (6 Gy) in A549 and H1299 cells by RT-qPCR using GAPDH as an endogenous control. Data were referred to unirradiated and untreated cells (control). Bars mean S.D. A549 (E) and H1299 (F) cells pretreated as in A and then exposed or not to navitoclax (1 μM) for 72 hours starting at day 5 after IR. Then media were replaced with navitoclax-free medium until the end of the experiment. Surviving fraction was normalized to respective unirradiated controls or unirradiated pretreated cells with navitoclax. Curves were fitted using lineal-quadratic model. Bars mean S.D.

Interestingly, our initial observation showed that in cells with null or dysfunctional p53, dinaciclib is unable to promote radiosensitivity, so we decided to test the role of p53 in the radiosensitizing effect of dinaciclib. To this end, we knocked down p53 expression in A549 cells using a specific shRNA. As shown, after effective knockdown of p53 (Figure 5A), the radiosensitising effect of dinaciclib was drastically reduced, correlating with a lack of senescence (Figure 5B, C and D). Collectively, these data support that p53-mediated radiation-associated senescence is an important determinant of the radiosensitising effect of dinaciclib.

**Fig. 5.**
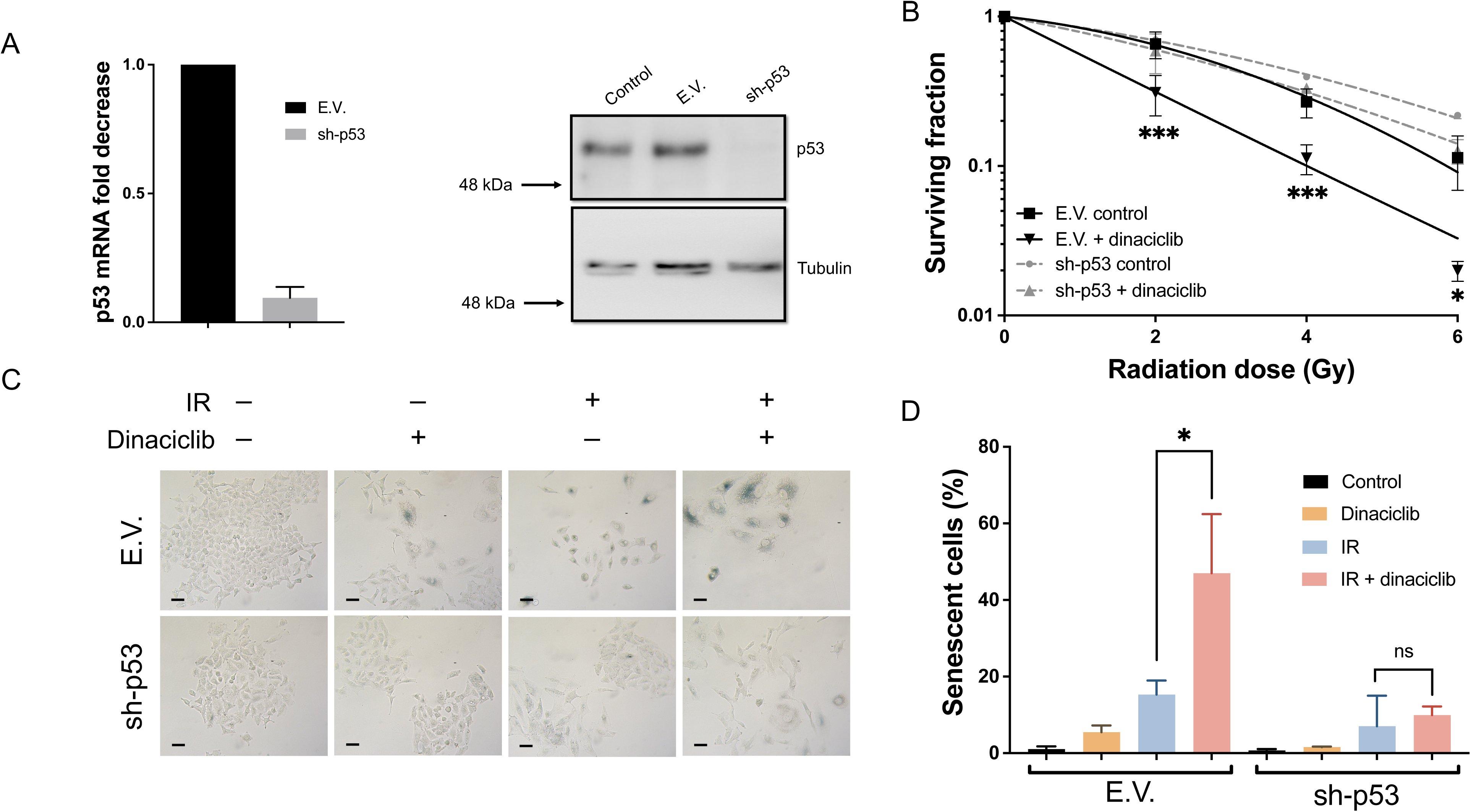
p53 is required for dinaciclib associated radiosensitivity. A) A549 cells were infected with lentiviruses carrying empty vector (E.V.) or shRNA for p53 (sh-p53). Selected pools were evaluated by RT-qPCR (left panel) and by western blot using tubulin as a loading control (right panel). Bars mean S.D. B) A549 cells infected with lentiviruses carrying E.V. or sh-p53 were plated and 24 hours later exposed to vehicle or dinaciclib (10 nM) for additional 24 hours. Then cells were exposed to the indicated doses of X rays and immediately after media was replaced.. Cellular radiosensitivity was plotted using respective control cells. Curves were fitted using lineal-quadratic model. Bars mean S.D. C): E.V (upper panels) or sh-p53 (lower panels) cells were irradiated (6 Gy) in presence/absence of dinaciclib 24 hours pretreatment (10 nM) and 6 days later β-Gal activity was detected by X-gal staining. A representative image is shown. Scale bars represent 100 μm. D) Histogram showing the average of, at least, three independent experiments representing the percentage of positive senescent cells. Bars mean S.D.

### 3.3. Dinaciclib blocks HR

In light of these findings, we next sought to investigate an upstream mechanism that controls the radiosensitive phenotype associated with dinaciclib. In this regard, abrogation of BRCA1 expression is one of the direct effects of dinaciclib [4]. As shown in Figure 6A, a significant decrease in BRCA1 expression, at both protein and RNA levels, was observed after exposure to dinaciclib in both models as well as in the other cell lines (Supplementary information 4). We therefore considered that the absence of BRCA1 could affect the DNA damage response and, more specifically to HR. To this end, we switched to an experimental model based on U2OS cells with functional p53 and HR [26]. As shown, dinaciclib was also able to promote radiosensitivity in U2OS cells (Figure 6B). Furthermore, in agreement with a decrease in BRCA1 expression, dinaciclib treatment showed a marked block of HR without affecting NHEJ, as measured by specific repair pathways reporters [23,24] (Figure 6C). To fully support the role of BRCA1, A549 cells were knocked down for BRCA1. After achieving effective knockdown (Supplementary information 5A), cells were exposed to IR in the presence/absence of dinaciclib pretreatment. As expected, the absence of BRCA1 almost block radiosensitivity associated to dinaciclib (Figure 6D). In fact, a marked senescence phenotype in response to IR was observed in cells lacking BRCA1 that remains unaffected by the dinaciclib pretreatment (Figure 6E and F). Furthermore, all these results were confirmed by a second shRNA (Supplementary information 5B, C and D).

**Fig. 6.**
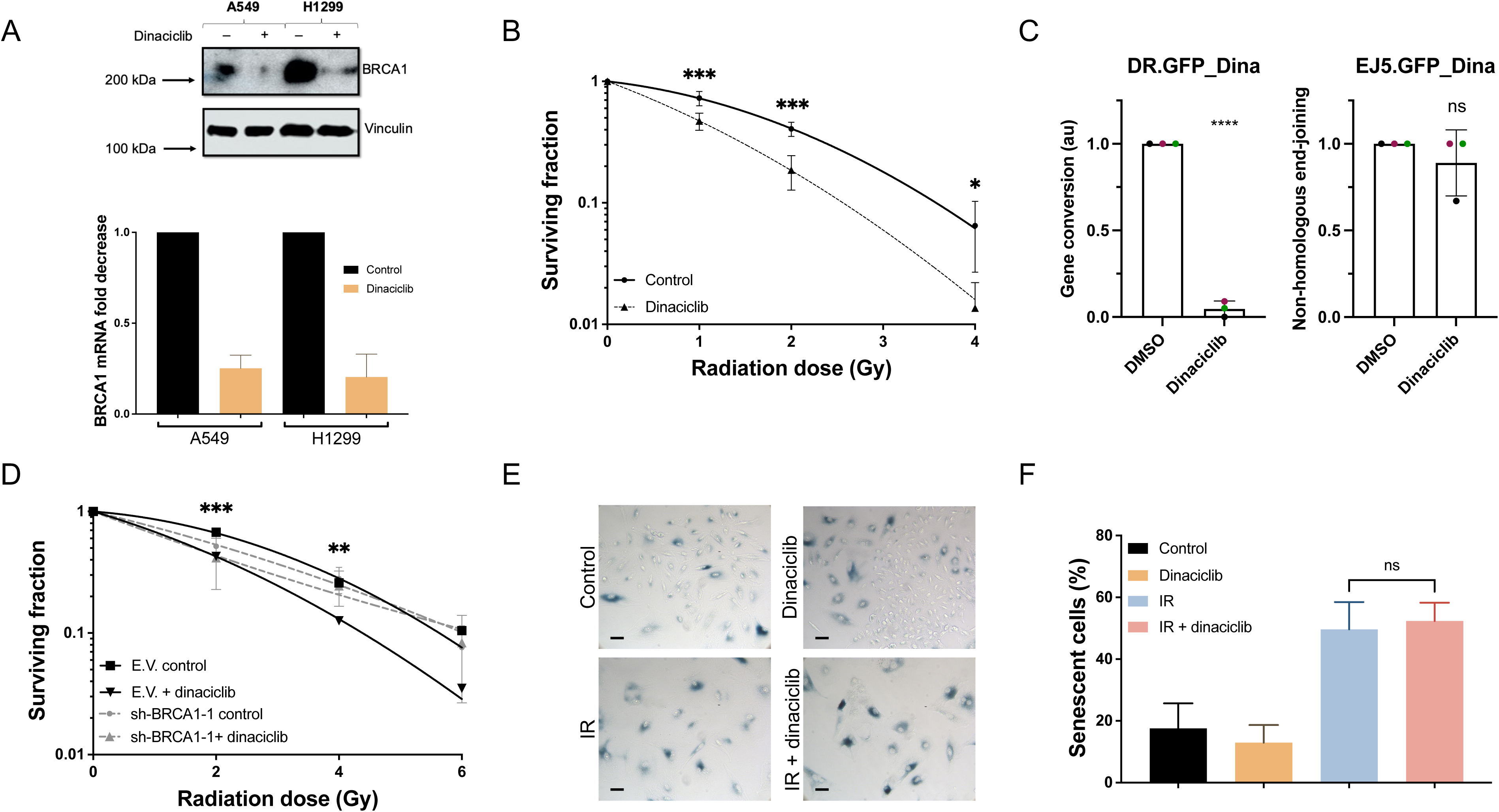
Downregulation of BRCA1 expression mediates radiosensitivity associated to dinaciclib. A) BRCA1 expression was evaluated in A549 and H1299 cell lines after 24 hours treatment of dinaciclib (10 nM) by western blot using vinculin as a loading control (upper panel) and by RT-qPCR (lower panel). B) Clonogenic assays in U20S cells. Cells were plated and 24 hours later exposed to vehicle or dinaciclib (10 nM) for additional 24 hours. Then cells were exposed to the indicated doses of X rays and immediately after media was replaced. Surviving fraction was normalized to respective unirradiated controls. Curves were fitted using lineal-quadratic model. Bars mean S.D. C) GFP reporter assay in U20S using DR.GFP (left histogram) or EJ5.GFP (right histogram). Histogram showing the average of three independent experiments. Bars mean S.D..D) A549 cells infected with lentiviruses carrying shRNA for BRCA1 (sh-BRCA1-1) were exposed to the indicated doses of X-rays in the presence of 24 hours pretreatment of dinaciclib (10 nM) or vehicle. Cellular radiosensitivity was plotted using control cells. Bars mean S.D. E) A549 cells infected with lentiviruses carrying sh-BRCA1 were irradiated (6 Gy) in presence/absence of 24 hours pretreatment of (10 nM) or vehicle and 6 days later SA-β-Gal activity was detected by X-Gal staining. A representative image is shown. Scale bars represent 100 μm. F) Histogram showing the average of, at least, three independent experiments representing the percentage of positive senescent cells. Bars mean S.D.

Finally, we decided to investigate the role of CDK12, which has been proposed as a key mediator of dinaciclib blockade of BRCA1 expression [4]. Therefore, silencing of CDK12 expression was performed using shRNA (Supplementary Information 6A). As shown, cells lacking CDK12 expression showed a marked downregulation of BRCA1 expression (Figure 7A). To fully elucidate the role of this particular CDK in dinaciclib-associated radiosensitivity, we performed a clonogenic assay in the presence/absence of dinaciclib pretreatment. Interestingly, dinaciclib was unable to promote radiosensitivity in CDK12-interfered cells (Figure 7B), which correlated with a marked senescent phenotype (Figure 7C and D). To fully support these observations, a second shRNA against CDK12 was challenged with almost identical results (Supplementary Information 6B, C and D).

**Fig. 7.**
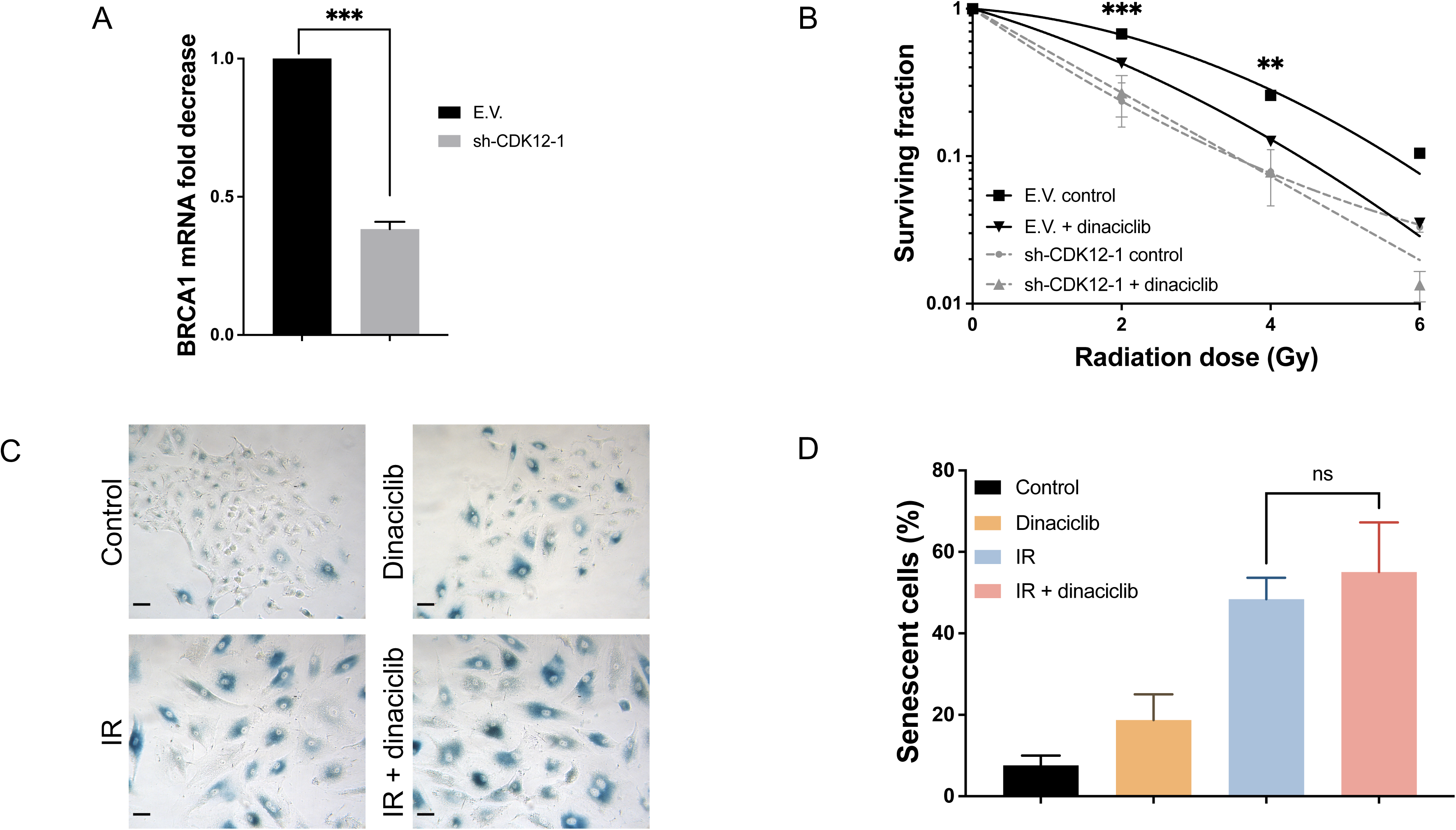
CDK12 inhibition is a key event dinaciclib associated radiosensitivity. A) BRCA1 expression was evaluated in A549 cells infected with lentiviruses carrying shRNA for CDK12 (sh-CDK12-1) by RT-qPCR (left panel) and by western blot using vinculin as a loading control (right panel). Bars mean S.D. B) A549 cells infected with lentiviruses carrying sh-CDK12-1 were exposed to the indicated doses of X-rays in the presence of 24 hours pretreatment of dinaciclib (10 nM) or vehicle. Cellular radiosensitivity was plotted using control cells. Bars mean S.D. C) A549 cells infected with lentiviruses carrying sh-CDK12-1 were irradiated (6 Gy) in presence of 24 hours pretreatment of dinaciclib (10 nM) or vehicle and 6 days later SA-β-Gal activity was detected by X-Gal staining. Representative images are shown. Scale bars represent 100 μm. D) Histogram showing the average of, at least, three independent experiments representing the percentage of positive senescent cells. Bars mean S.D.

Therefore, these results support a model in which CDK12 inhibition abrogates BRCA1 expression, rendering a nonfunctional HR that increases senescence associated to IR in a p53 dependent manner which mediates radiosensitivity associated to dinaciclib.

## 4. Discussion

Several conclusions are drawn from the present report.

First, dinaciclib is a novel and potent radiosensitiser that could be added to the growing list of cell cycle-targeted therapies with the ability to induce radiosensitivity [27,28], but it cannot be claimed to be a universal radiosensitiser. In this sense, our data allow the correct selection of patients who could benefit from a future combination therapy based on markers such as BRCA1 or p53, which are currently being evaluated in clinical practice. This observation could be of particular interest in pathologies such as breast or lung cancer, where radiotherapy is a cornerstone and dinaciclib is being considered for clinical use [6,7,29,30].

Second, dinaciclib promotes radiosensitivity in a genetic context that requires at least functional p53 and BRCA1. Regarding p53, it is important to mention that p53 is one of the most common alterations in cancer present in approximately 50% of tumours [31], suggesting that our finding could be limited to a group of tumours with a theoretically better response to IR. However, several points highlight the importance of the current findings. First, the definitive role of p53 in cellular radioresistance is not clear [32]. Furthermore, no clinical protocol considers p53 status as a biomarker for radiotherapy. For example, in silico searches in The Cancer Genome Atlas Database (TCGA) lung squamous cell carcinoma cohort using the cBioPortal tool [33], showed that WT p53 tumours did not have a better prognosis than mutant p53 tumours, regardless of whether they were treated with radiotherapy or not (Supplementary Information 7). Therefore, there is an urgent clinical need to optimize radiotherapy not only for p53 mutant tumours but also for WT p53 tumours, reinforcing the importance of our findings. Our observations on the role of p53, obtained using different models such as HT-29 or H1299 as well as shRNA experiments, showed results along the same lines as those obtained with palbociclib [34] or abemaciclib [35]. However, it is known that both CDK4/6 inhibitors directly affect ATM kinase activity and subsequent activation of p53 [34,36], whereas dinaciclib had no effect on ATM signalling as assessed by H2AX or KAP phosphorylation. Notably, recent studies indicate that dinaciclib is able to inhibit ATM/CHK2 signalling in Hela and SiHa cells, suggesting that cell cycle and apoptosis are the main mechanisms of dinaciclib-associated radiosensitivity [37]. However, despite the different experimental models used, these conclusions are based on a correlation that lacks genetic approaches and quantitative methods. In addition, it is noteworthy that in their two cellular models, IR is unable to activate CHK2 or H2AX phosphorylation in response to IR, which is totally opposite to our model as well as previous work in Hela cells showing normal activation of H2AX or CHK2 after IR [38,39]. Furthermore, in terms of apoptosis the results are similar to our study showing no synergistic effect, which clearly exclude apoptosis as a radiosensitizer mechanism. Finally, the cell cycle alteration shown by Zang and coworkers does not indicate whether dinaciclib is affecting G2/M exit after IR exposure and, interestingly, no increase in G2/M is observed after IR exposure in any of the cell lines, which is totally unexpected, and opposite to what we have shown in our experimental setting as well as previous works in Hela cells [40]. In conclusion, this previous work needs to be carefully re-evaluated to avoid misinterpretation in order to optimise this novel combination.

Third, the effect of dinaciclib on BRCA1 expression is a key issue. Indeed, it has been shown that cells lacking BRCA1 are extremely sensitive to IR [41] and prone to senescence [42] mediated by p53 [43], fitting perfectly with our proposed model of dinaciclib-associated radiosensitivity. In this sense, dinaciclib could be acting in a similar way to other HR blockers as tanespimycin [44], a known inhibitor of Hsp90 that blocks HR [45] and confer radiosensitivity by increasing p53-dependent senescence through its effect on BRCA1 proteasomal degradation [46,47]. Furthermore, in an attempt to bring our observation to the clinical level, we performed in silico studies (Supplementary information 8) about the level of BRCA1 expression in the TCGA lung squamous cell carcinoma cohort. Interestingly, we did not observe correlation with overall survival (P-value: 0.487). However, if we analyse the population that received radiotherapy, 52 out of 486 patients, we observe that BRCA1 expression levels almost reach statistical significance (Supplementary information 8B, P-value: 0.0532), suggesting that high levels of BRCA1 may mediate a poor response to radiotherapy. Although these data may support our hypothesis, further studies are needed to draw more definitive conclusions at the clinical level as has been proposed for BRCA1 mutations [48]. Nonetheless other genes regulated by dinaciclib as Rad51 [4] should be considered which are also critical in the HR process [49], and known to control radiosensitivity [50,51].

In addition, our data suggest that senescence is the critical biological process for the radiosensitising effect associated with dinaciclib, providing the first link between radiation-induced senescence and dinaciclib. This observation opens the possibility of considering senolytics as a new player in our proposed chemo/radiotherapy regimen. However, it is important to note that the pro-senescent effect of dinaciclib could be stimulus-dependent, as has been shown in the case of doxorubicin [11] but not for arginine-deprivation [52]. Therefore, our data support the possibility of a therapeutic regimen based on dinaciclib/radiotherapy/senolytics in a sequential manner, which could significantly improve the potential of radiotherapy [53,54]. Furthermore, our observation may have implications in others therapeutic approaches as immunotherapy in which senescence has recently been shown to be involved [55] and interestingly could explain the effect observed for the combination of dinaciclib and immunotherapy based on anti-PD-1 [56].

Finally, another interesting question raised by our data is the role of CDK12 in the combination of dinaciclib and radiotherapy. Our data support the role of CDK12 onto BRCA1 expression as the critical step to explain the effect of dinaciclib in response to IR. In fact, our findings open the possibility to consider CDK12 inhibitors not only as new targeted therapy agents [57,58] but also as potent radiosensitising agents trough the modulation of radiation induced senescence. However, we cannot rule out other dinaciclib targets as CDK9, that has been reported to control a BRCA1 recruitment to DNA damage sites [59] and also expression levels [60].

In conclusion, we present new evidence that dinaciclib is a potent radiosensitising agent through its inhibitory effect on CDK12, which downregulates the expression of key components of the HR machinery such as BRCA1. This event promotes a senescent phenotype that requires a WT p53. Our observations provide the rationale for a future combination of radiotherapy plus dinaciclib in those patients with a functional BRCA1/p53 axis, which accounts for approximately 50% of cases, allowing for more selective and effective radiotherapy.

## Abbreviations

(CDKs): Cyclin-dependent kinases
(IR): Ionising radiation
(SA-β-Gal): Senescence-Associated β-Galactosidase
(HR): Homologous Recombination
(NHEJ): Non-Homologous End Joining
(shRNA): Short hairpin RNA
(TGCA): The Cancer Genome Atlas
(S.D.): Standard deviation
(SF): Surviving fraction
(SASP): Senescence-associated secretory phenotype
(WT): Wild type

