## Supplementary material for "The signalling axis CDK12-BRCA1 mediates dinaciclib associated radiosensitivity through p53-mediated cellular senescence": Supplemnetary files and tables

Supl.1

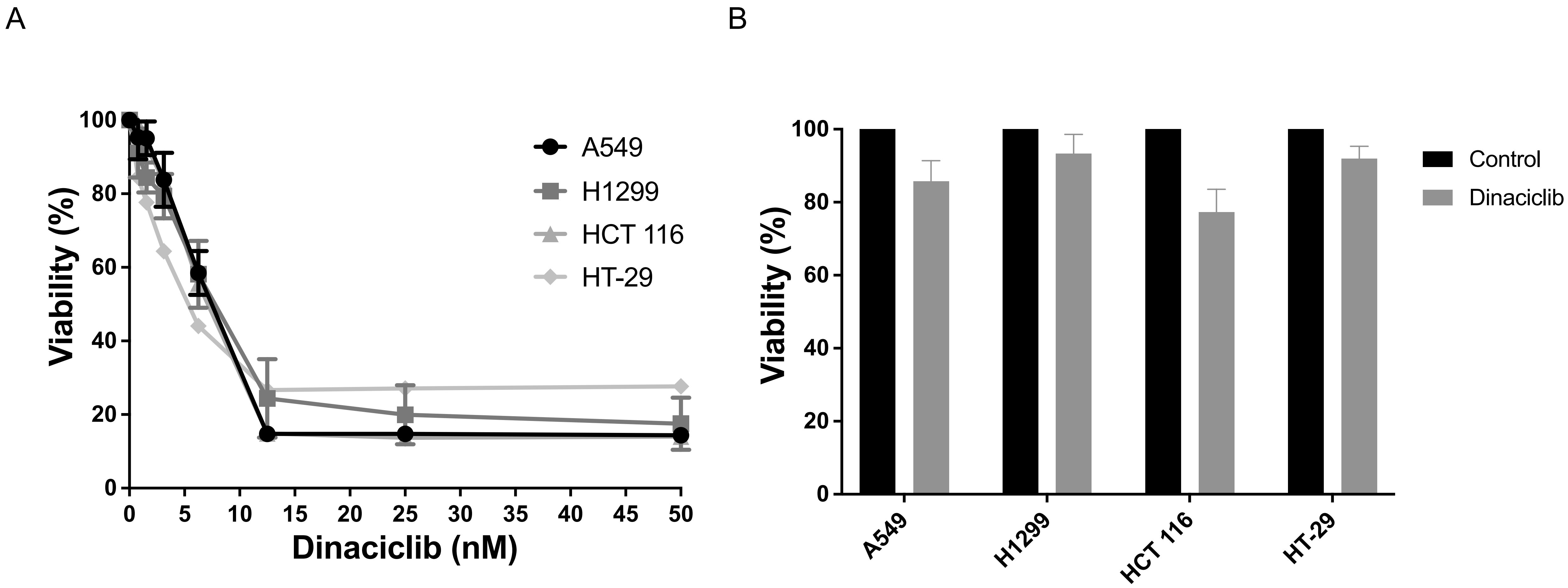


A) Cells were plated in a 24 well plate (1x10⁴ cells/well) 16 to 24 hours prior to treatment with dinaciclib (doses ranging from 0 to 50 nM) during 72 hours, viability was assayed by MTT assay. B) Same as in A but using vehicle or dinaciclib 10 nM for 24 hours.

Supl. 2

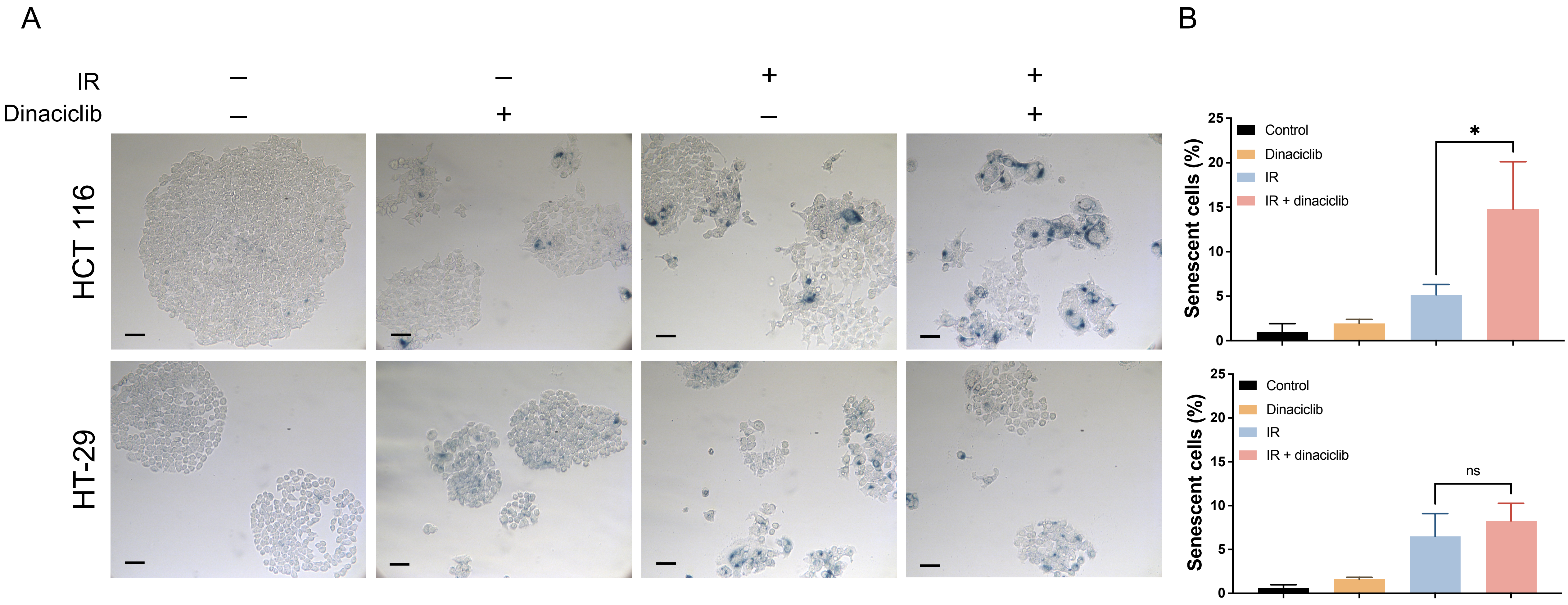


A) HCT 116 (upper panels) and HT-29 (lower panels) cells pretreated with dinaciclib (10 nM) or vehicle for 24 hours were irradiated (6 Gy) and 6 days later SA-β-Gal activity was detected by X-Gal staining. A representative image is shown for each cell line. Scale bars represent 100 μm. B) Histogram showing the average of, at least, three independent experiments. Bars mean S.D. HCT 116 (upper panel) and HT-29 (lower panel).

Supl. 3

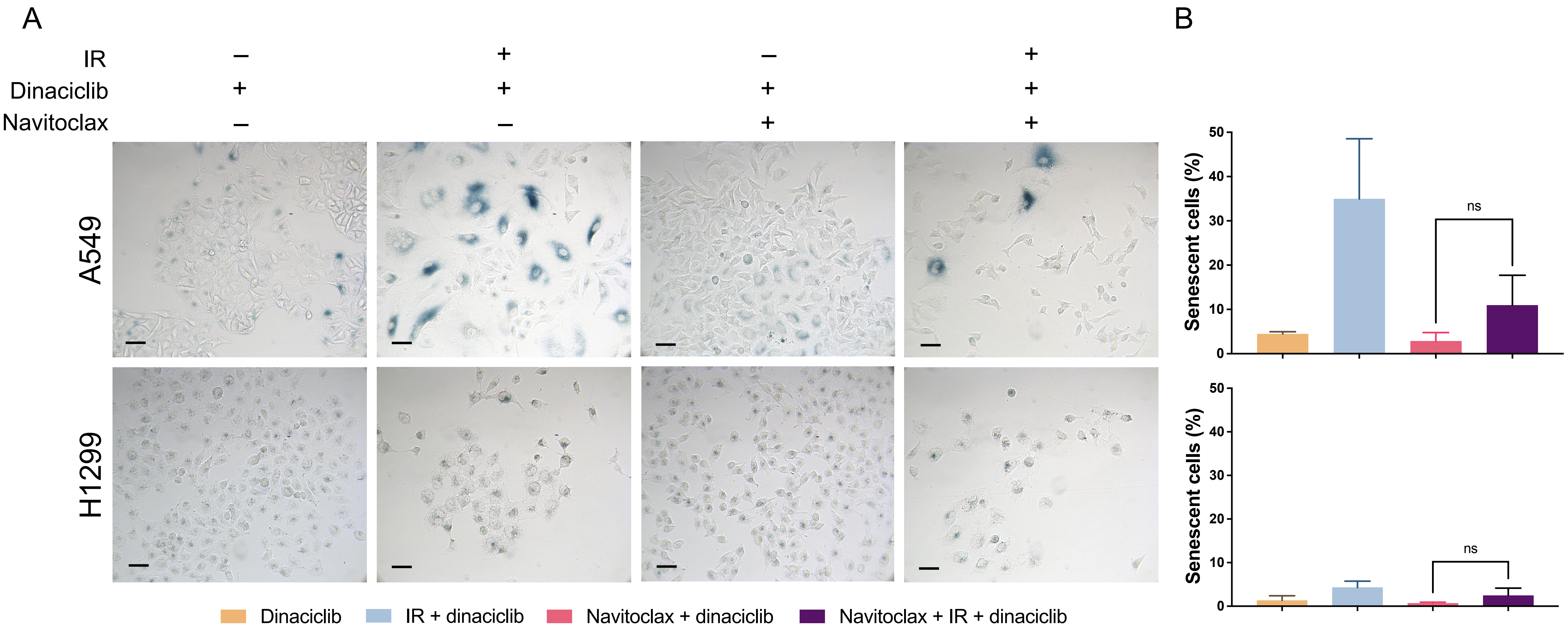


A) A549 (upper panels) and H1299 (lower panels) cells pretreated with dinaciclib (10 nM) for 24 hours were irradiated (6 Gy) and 5 days after IR exposure, cells were treated with navitoclax (1 μM) for 72 hours and then, SA-β-Gal activity was detected by X-Gal staining. A representative image is shown for each cell line. Scale bars represent 100 μm. B) Histogram showing the average of, at least, three independent experiments. Bars mean S.D. A549 (upper panel) and H1299 (lower panel).

Supl. 4

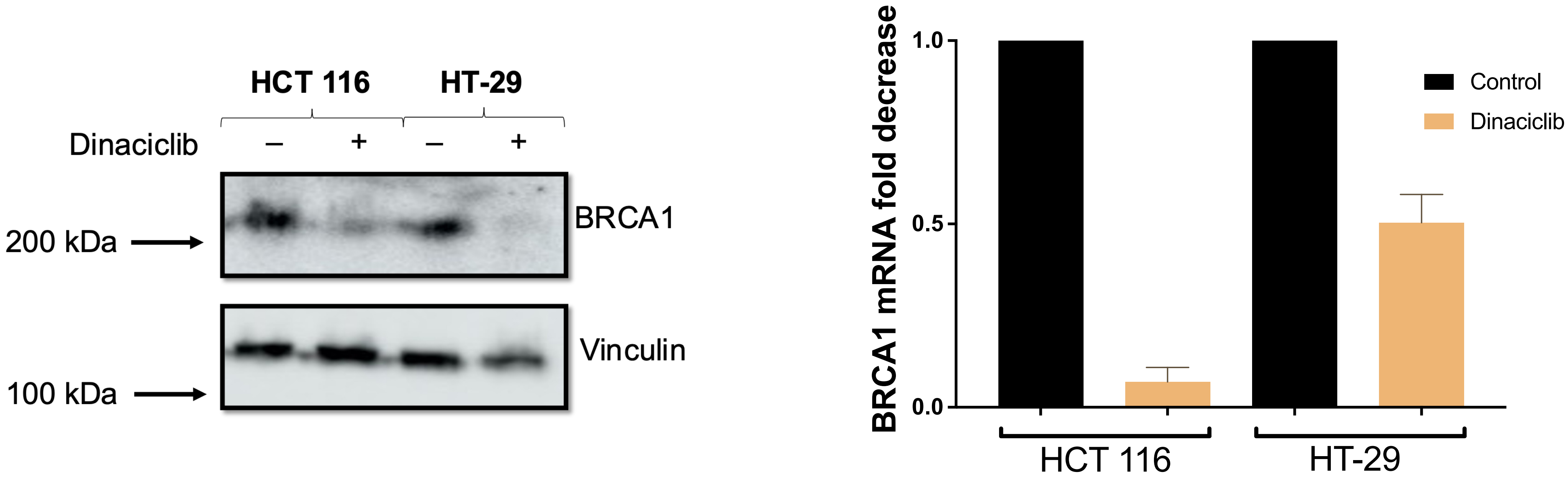


BRCA-1 expression was evaluated in HCT 116 and HT-29 cell lines after 24 hours treatment of dinaciclib (10 nM) by western blot using vinculin as a loading control (left panel) and by RT-qPCR using GAPDH as an endogenous control (right panel).

Supl. 5

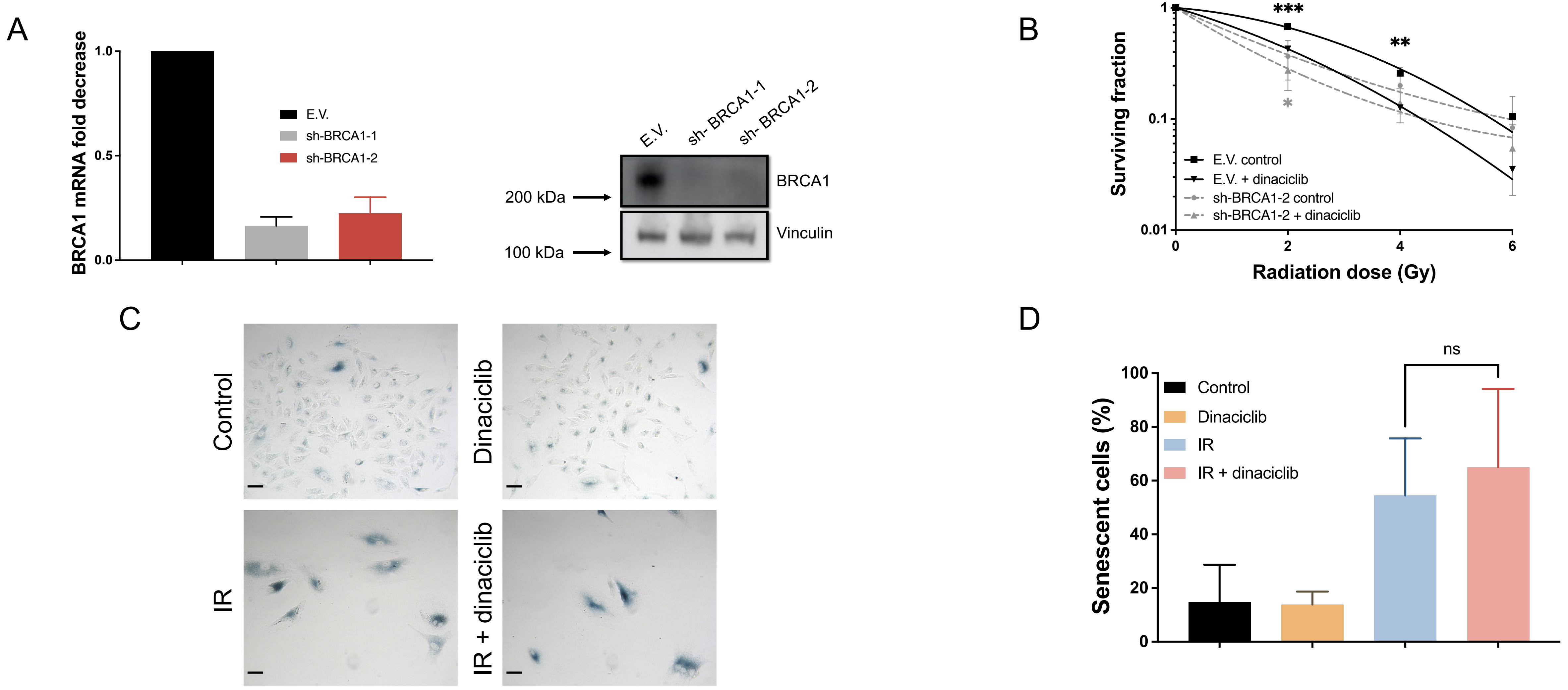


A) BRCA1 expression was evaluated in A549 cells infected with lentiviruses carrying shRNAs for BRCA1 (sh-BRCA-1 and sh-BRCA1-2) by RT-qPCR (left panel) and by western blot using vinculin as a loading control (right panel). B) A549 cells infected with lentiviruses carrying sh-BRCA1-2 were exposed to the indicated doses of X-rays in the presence of 24 hours pretreatment of dinaciclib (10 nM) or vehicle as in figure 6D. C) A549 cells infected with lentiviruses carrying sh-BRCA1-2 were treated and processed as in figure 6E. Representative image is shown. Scale bars represent 100 µm. D) Histogram showing the average of, at least, three independent experiments representing the percentage of positive senescent cells.

Supl 6

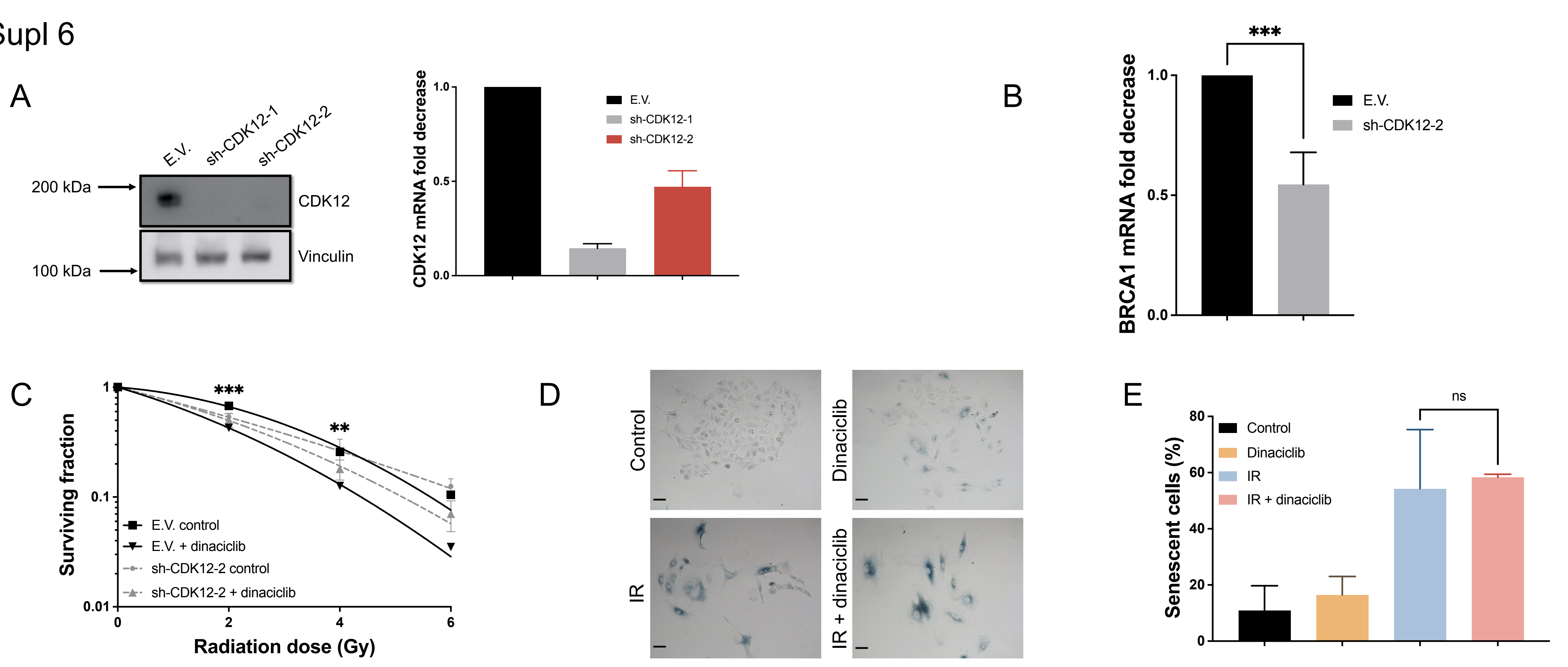


A) CDK12 expression was evaluated in A549 cells infected with lentiviruses carrying shRNAs for CDK12 (sh-CDK12-1 and sh-CDK12-2) by RT-qPCR (left panel) and by western blot (right panel). B) BRCA1 expression was evaluated in A549 cells infected with lentiviruses carrying sh-CDK12-2 by RT-qPCR (left panel) and by western blot using vinculin as a loading control (right panel). C) A549 cells infected with lentiviruses carrying sh-CDK12-2 were exposed to the indicated doses of X-rays in the presence of 24 hours pretreatment of dinaciclib (10 nM) or vehicle as in figure 7B. D) A549 cells infected with lentiviruses carrying sh-CDK12-2 were treated and processed as in figure 6E. Representative image is shown. Scale bars represent 100 µm. E) Histogram showing the average of, at least, three independent experiments representing the percentage of positive senescent cells.

Supl. 7

A

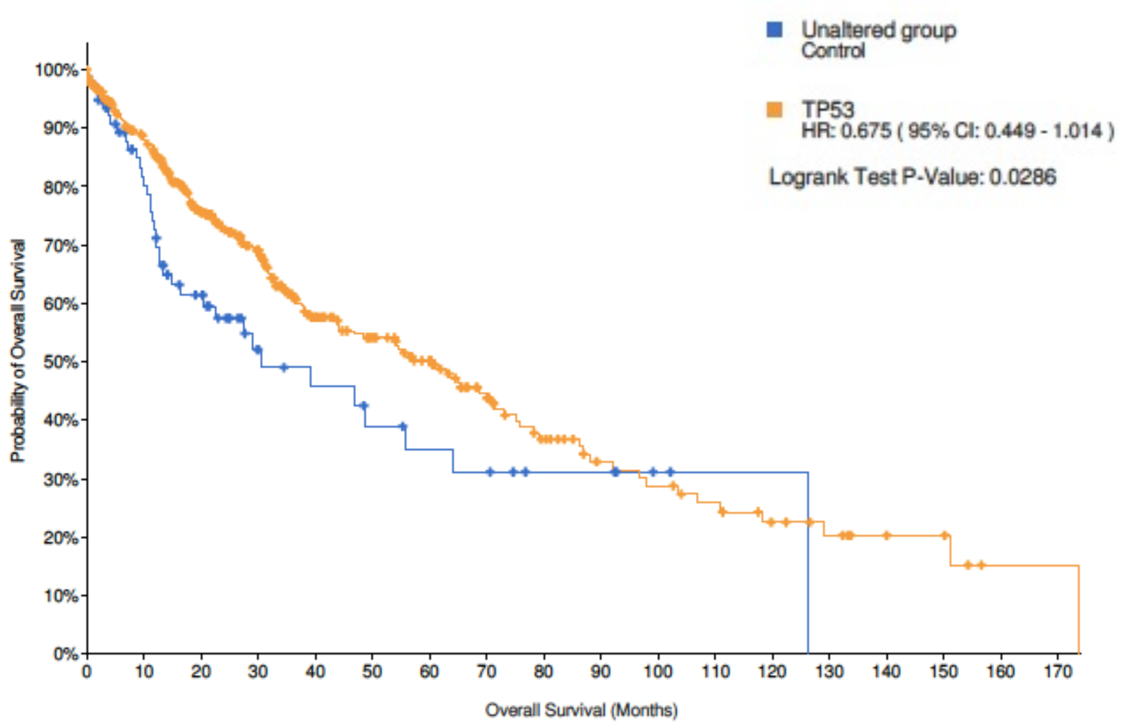


**Number at risk (n)**

| | | | | | | | | | | | | | | | | | | |
|---|---|---|---|---|---|---|---|---|---|---|---|---|---|---|---|---|---|---|
| Unaltered group | 79 | 53 | 34 | 17 | 14 | 11 | 9 | 8 | 5 | 5 | 2 | 1 | 1 | 0 | 0 | 0 | 0 | 0 |
| TP53 | 399 | 309 | 220 | 166 | 110 | 88 | 71 | 50 | 34 | 24 | 21 | 17 | 12 | 9 | 6 | 5 | 1 | 1 |

B

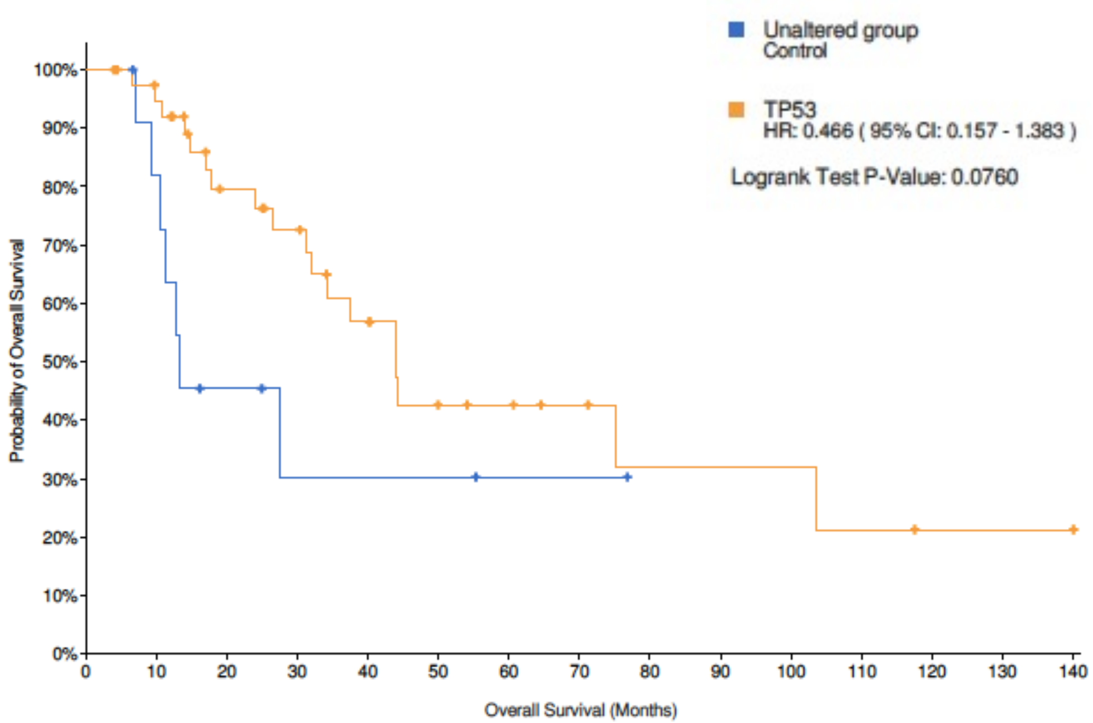


**Number at risk (n)**

| | | | | | | | | | | | | | | | |
|---|---|---|---|---|---|---|---|---|---|---|---|---|---|---|---|
| Unaltered group | 12 | 9 | 4 | 2 | 2 | 2 | 1 | 1 | 0 | 0 | 0 | 0 | 0 | 0 | 0 |
| TP53 | 40 | 35 | 24 | 20 | 14 | 8 | 7 | 5 | 3 | 3 | 3 | 2 | 1 | 1 | 1 |

A) Kaplan–Meyer comparing prognosis in terms of Overall Survival for two groups of patients, those with mutant p53 (yellow line) and WT (blue line) in the lung squamous cell carcinoma cohort from TCGA dataset (487 patients). Statistical significance of the difference was evaluated by logrank test.
B) Same as in A but for patients that were irradiated (52 patients).

Supl. 8

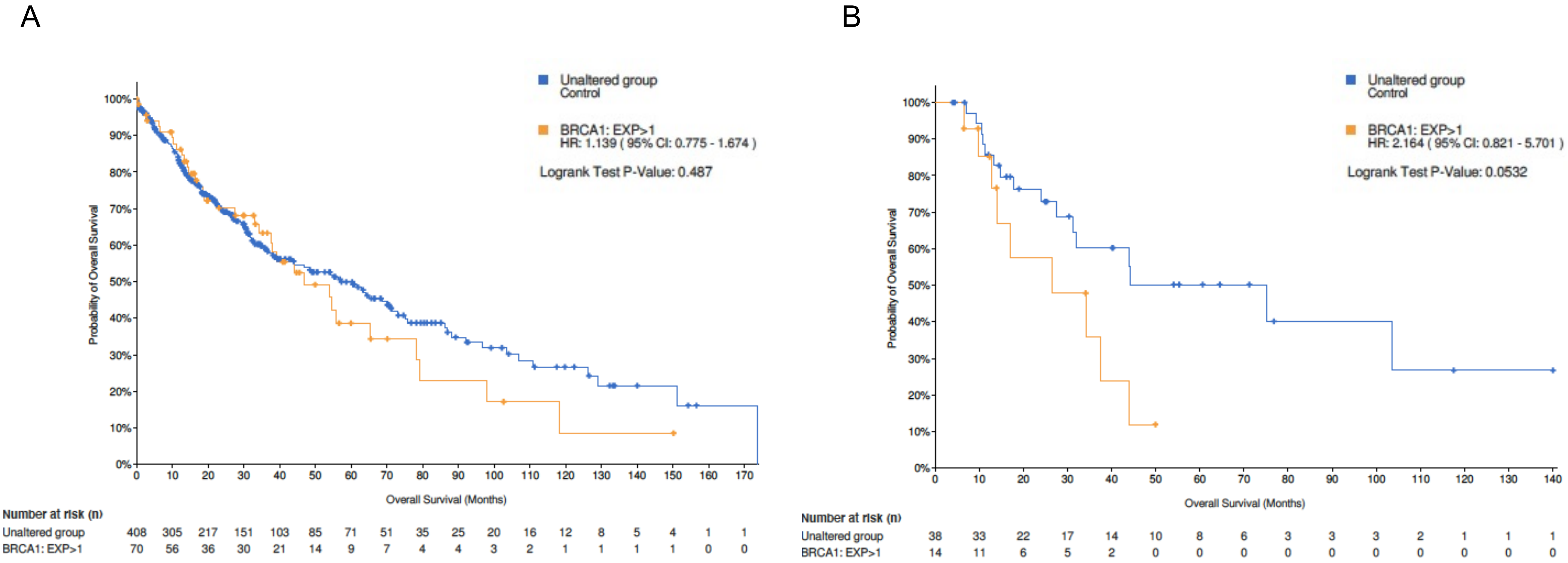


A) Kaplan–Meyer comparing prognosis in terms of Overall Survival for two groups of patients, those with high (mRNA expression > 1 S.D) and low (mRNA expression < 1. S.D) expression levels of BRCA1 in the lung squamous cell carcinoma cohort from TCGA dataset (487 patients). Statistical significance of the difference was evaluated by logrank test. B) Same as in A but for patients that were irradiated (52 patients).

Supl. table 1

| Use | Antibody | Dilution | Manufacturer | Reference |
|---|---|---|---|---|
| Immunocytochemistry | p-H2AX | 1:500 | Cell Signaling Technology | 9718 |
| Immunocytochemistry | Anti-rabbit Alexa Fluor 488 | 1:2000 | Invitrogen | A-11008 |
| Western Blot | P53 | 1:500 | Santa Cruz Biotechnology | sc-47698 |
| Western Blot | BRCA1 | 1:1000 | Cell Signaling Technology | 9010 |
| Western Blot | KAP1 | 1:1000 | Bethyl Laboratories | A300-274A-T |
| Western Blot | p-KAP1 | 1:1000 | Bethyl Laboratories | A700-013-T |
| Western Blot | CDK12 | 1:1000 | Cell Signaling Technology | 11973 |
| Western Blot | Tubulin | 1:2000 | Santa Cruz Biotechnology | sc-32293 |
| Western Blot | Vinculin | 1:3000 | Sigma-Aldrich | V9264 |
| Western Blot | Anti-mouse-HRP | 1:2000 | Cell Signaling Technology | 7076 |
| Western Blot | Anti-rabbit-HRP | 1:2000 | Cell Signaling Technology | 7074 |

Supl. table 2

| **Use** | **Target** | **Forward** | **Reverse** |
|---|---|---|---|
| RT-qPCR | *GAPDH* | TCGTGGAAGGACTCATGACCA | CAGTCTTCTGGGTGGCAGTGA |
| RT-qPCR | *IL-1β* | TGCACGTCCGGGACTCACA | CATGGAGAACACCACTTGTTGCTCC |
| RT-qPCR | *IL-6* | GATGAGTACAAAAGTCCTGATCC | CTGCAGCCACTGGTTCTGT |
| RT-qPCR | *IL-8* | AGACAGCAGAGCACACAAGC | ATGGTTCCTTCCGGTGGT |
| RT-qPCR | *BRCA1* | TAGGGCTGGAAGCACAGAGT | AATTTCCTCCCCAATGTTCC |
| RT-qPCR | *CDK12* | TGAAAACCCAAGAGCCAGCA | GTGGAAGAATGTGAGGAGGACAT |
| RT-qPCR | *TP53* | TCCTGCCATTTTGGGTTT | GCAGGCCAACTTGTTCAGTG |
